# Symbiont effectors modulate plant signalling to increase host stress resilience

**DOI:** 10.64898/2026.08.17.744369

**Authors:** Laura Rehneke, Rory Osborne, Silke Lehmann, Yingqi Zhang, Jemma Roberts, Stefan Altmann, Eva Köpff, Ruth Eichmann, Pascal Falter-Braun, Weixing Shan, Patrick Schäfer

## Abstract

Plant colonizing mutualistic symbionts confer beneficial effects to their hosts, which often includes increased growth, and biotic and abiotic stress resilience. How these benefits are activated on a molecular level is mostly unknown. Here, we describe effector candidates of the fungal symbiont *Serendipita indica* (*Si*), which modulate plant stress signalling pathways. By analyzing the *Si* effector interactome, we reveal frequent targeting of stress related host proteins, which are linked to the identified effector signalling functions. Moreover, functional data indicate that *Si* effectors modulate abiotic stress response of Arabidopsis, as well as resistance to pathogen infection. Analysis of symbiont effectors might not only uncover previously unreported molecular mechanisms that increase plant fitness but might also be used to identify potential genetic traits for crop improvement under changing climates.

## Introduction

As sessile organisms, plants cannot escape environmental fluctuations and need to adapt to stress events. These stresses can be abiotic, including drought, cold, heat, high light and salinity, or biotic stresses caused by pathogens and insects. The molecular responses of plants to these stresses are supported by a highly interconnected hormone signalling network. In response to unfavourable conditions plants activate extensive hormone signalling for adaptation and defence (Verma et al. 2016; Hou and Tsuda 2022; Ku et al. 2018). Strongly interconnected hormone signalling pathways regulate all aspects of plant life (Hou and Tsuda 2022; Emenecker and Strader 2020; Khan et al. 2020; Vega et al. 2019; Altmann et al. 2020). As a result, increased crop stress resistance is linked to essential plant processes and particularly to growth-related pathways. To alleviate the impact of climate change on global agriculture and the growing number of people falling into food insecurity (Ahmadalipour et al. 2019; Mirón et al. 2023), it will be necessary to identify measures for plants to manage growing environmental stresses to reduce overall crop losses.

Plants have evolved a complex network of hormones to integrate developmental and environmental cues. These include abscisic acid (ABA) and auxin (AUX) which are canonically associated with growth and development (Emenecker and Strader 2020), and salicylic acid (SA) and jasmonic acid (JA) which are widely investigated in plant immunity (Hou and Tsuda 2022). Due to the complexity of the hormone network, no plant hormone operates in isolation, with many points of crosstalk having evolved within the network to co-regulate different processes and stress adaptation (Ku et al. 2018; Yang et al. 2019; Altmann et al. 2020; García-Andrade et al. 2020).

Beneficial symbionts are part of microbial communities and have been shown to enhance crop fitness under many stress conditions, including cold (Acuña-Rodríguez et al. 2020), heat, salinity, drought and extreme temperatures (Boorboori and Zhang 2022; Mitra et al. 2021; Gupta et al. 2021; Begum et al. 2019; Latef et al. 2016; Harman 2011). At the same time beneficial microbes also improve defence mechanisms against pathogens. Induced systemic resistance (ISR) is a priming process used by growth-promoting bacteria and fungi to enhance host defence against pathogens and herbivores (Pieterse et al. 2014). Beneficial symbioses are typically associated with improved nutrient exchange. Mutualists are supplied with carbon and in return supply their host with essential minerals, with many terrestrial plants depending on beneficial fungi for nutrient supply (Acuña-Rodríguez et al. 2020; Behie and Bidochka 2014). Improved nutrient subsistence, however, cannot explain the diversity of beneficial properties symbiont-colonized host plants acquire (Yan et al. 2019; Xu et al. 2018), suggesting additional mechanisms are employed by symbiotic microbes to promote beneficial effects in host plants.

The symbiont *Serendipita indica* (*Si*) stimulates growth and development in addition to increasing resilience to both biotic and abiotic stresses (Baltruschat et al. 2008; Vadassery and Oelmüller 2009). So far, these effects have been found to be associated with the ability of the symbiont to influence plant ROS levels, but also the manipulation of host hormone networks (Xu et al. 2018; Nath et al. 2016). Similar to pathogens, symbionts must suppress host immunity (Jacobs et al. 2011) and utilize effectors to manipulate host immune responses in the course of colonization (Yang et al. 2022; Daneshkhah et al. 2018; Wawra et al. 2016; Jacobs et al. 2011). While microbes often target hormone homeostasis (Pozo et al. 2015), it is unclear to what extent effectors are involved and reprogram underlying signalling networks. Comparative interactome studies have revealed that *Si* effector candidates (SIECs) specifically target the Arabidopsis hormone network, and to a broader extent than the effector repertoires of different pathogens (Akum et al. 2015; Shen et al. 2018; Osborne et al. 2023). Further functional characterization of these putative effectors has demonstrated that those targeting hormone pathways can improve plant growth when expressed constitutively, highlighting their potential as tools to understand beneficial host signalling and symbiosis.

Here, we explore the beneficial properties of SIECs in plants exposed to abiotic and biotic stresses. Functional analyses in protoplasts revealed modulation of abiotic stress and pattern-triggered immunity (PTI) signalling. Integration of this information with previously published SIEC-Arabidopsis protein interactome revealed a diversity of SIECs which might co-ordinate stress responses in whole plants by altering target activity. In both directed and unbiased phenotyping assays, we demonstrate that SIECs increase host growth, improve resilience to abiotic stresses, and alter host susceptibility to pathogen infection.

## Results

### SIECs modulate stress and hormone signalling in protoplasts

We previously identified *S. indica* (*Si*) effector candidates (SIECs) involved in hormone signalling and root growth promotion (Osborne et al. 2023). Since hormones are required in major signalling and stress pathways of plants (Khan et al. 2020; Ku et al. 2018), we aimed to investigate the function of effector candidates in the modulation of stress responses. We previously analysed specific hormone signalling markers (pHORMONE) in *Arabidopsis thaliana* (Arabidopsis). Here we utilized *35S::SIEC* co-expressed with markers containing promotors (*pSTRESS*) of genes responsive to abiotic stress/heat (*DREB2a* and *HSP81.1*) or biotic stress/flg22-induced PTI (*PHI1* and *FRK1*) (Figure 1A). In a first landmark screen, 106 SIECs were tested against each marker in protoplasts (848 assays in total) (Figure 1A). 82 of 106 SIECs showed modulation of one or multiple stress markers. 49 and 40 SIECs altered expression of the abiotic stress markers *pDREB2a::LUC* and *pHSP81.1::LUC*, respectively. For PTI, 28 SIECs altered *pPHI1::LUC* and 26 *pFRK1::LUC* signalling (Figure 1B, Supplementary Table 1). The majority of SIECs only affected one or two pathways (Figure 1C) and most of these SIECs modulated only abiotic stress markers, 16 altered only PTI responses, and 28 affected both (Figure 1D). We validated the landmark screen by performing 3 additional biological replicates of the top 10 inducers and repressors of the *pDREB2a::LUC* and *pPHI1::LUC* markers in protoplast assays. Of the 20 respective validations performed, we were able to confirm effector function in 60% of cases by statistical verification of mock and treatment response relative to the empty vector control (Figure 1E, Supplementary Table 1). While somewhat low due to technical variations between biological repeats, this figure was still encouraging that effectors might modulate respective pathways in plants. Subsequent phenotyping assays revealed stress response alterations by effectors, which correspond to the signalling pathway function in protoplast assays (Figures 4F-I). Taken together, these protoplast assays indicated that *Si* utilizes effector proteins not only to alter immunity signalling, but also to modulate environmental stress responses.

**Figure 1.**
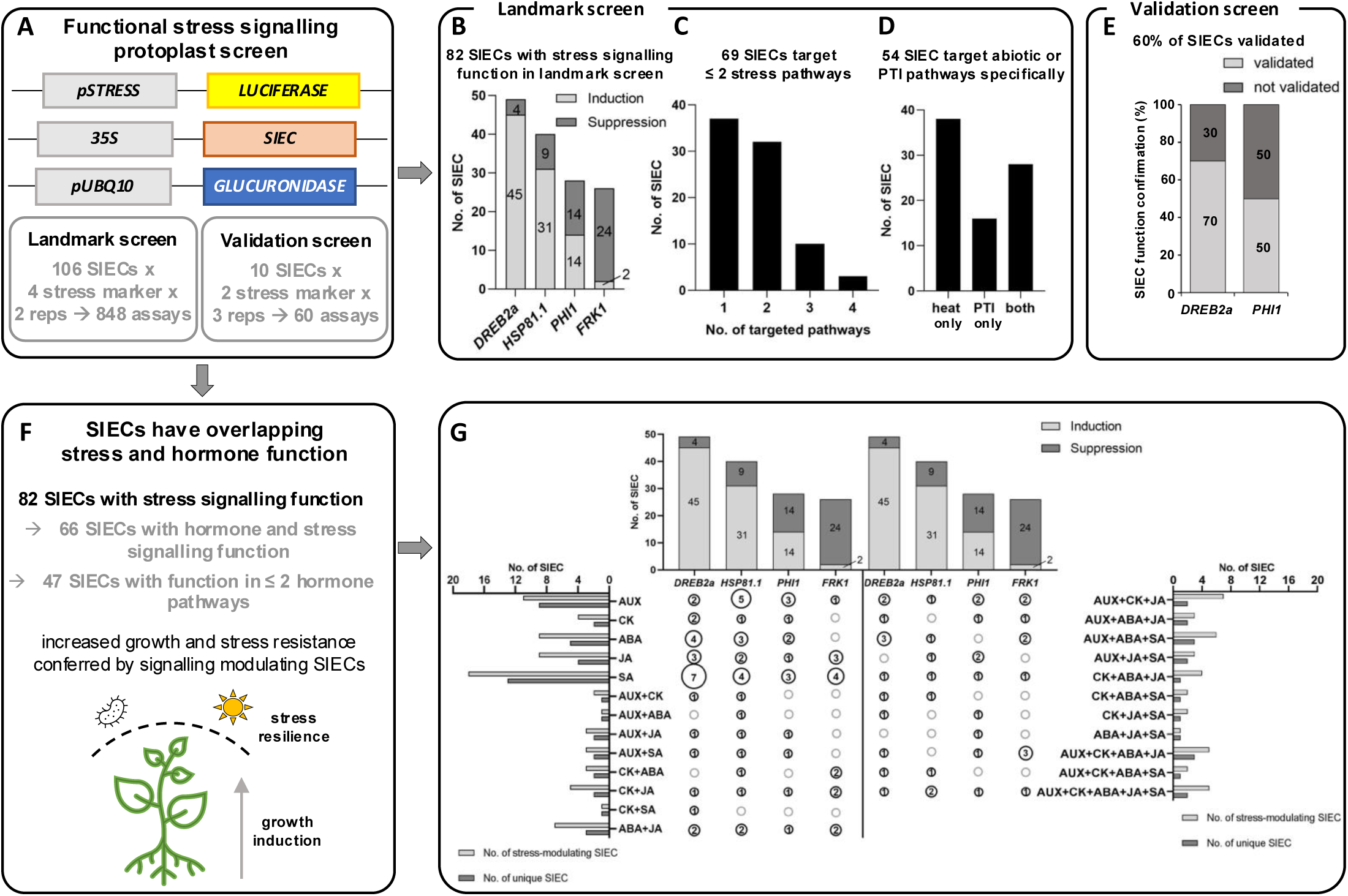
SIECs modulate stress and hormone signalling pathways. SIEC function on abiotic stress and immunity marker expression in protoplasts. **A** Overview of initial stress marker screen in protoplasts. **B** Number of SIECs inducing or repressing abiotic stress (*DREB2a* and *HSP81.1*) or PTI marker (*PHI1* and *FRK1*) expression calculated from multiplied ratios of mock and treatment conditions. **C** Number of SIECs modulating one or multiple marker pathways indicating marker-specific SIEC functions. **D** Number of SIEC modulating one or both stress signalling pathways indicating pathway-specific SIEC functions. **E** Confirmation of SIEC function in modulating *pDREB2a::LUC* and *pPHI1::LUC* markers, respectively. **F** Summary of SIECs with overlapping stress and hormone signalling modulating functions. **G** Number of SIECs with overlapping hormone and stress functions. Top bar chart shows number of SIECs modulating stress markers. Bar charts on the left and right indicate the number of SIECs additionally altering individual or multiple hormone markers (hormone data adapted from Osborne et al. 2023). Number of SIECs affecting different combinations of hormone markers and the individual stress pathways is listed between the different bar charts. AUX – auxin, CK – cytokinin, ABA – abscisic acid, JA – jasmonic acid, SA – salicylic acid.

Given that stress responses are tightly linked to plant hormone regulation, we combined our published hormone data (Osborne et al. 2023) with the present stress marker signalling data in an UpSet plot (Figure 1F, G). 66 SIECs modulate both hormone and stress signalling in protoplasts (80%, 66 out of 82 SIEC with stress function). In concordance with our previous observation that SIECs are highly specific in the hormone pathways they target, most effectors that modulated stress signalling affected only one hormone pathway (33 SIECs, Figure 1F, G). To investigate whether the hormone and stress function of SIECs was interconnected, we analysed the dataset toward possible groupings. SIECs only modulating immunity markers similarly affected all hormone markers (Supplementary Figure 1A). SIECs only altering heat stress response, also changed AUX, SA and ABA and to a lesser extent CK and JA regulation (Supplementary Figure 1A). While most SIECs that affected *pPHI1* and *pFRK1* activity also modulated JA signalling (Supplementary Figure 1B), the alignment of defence hormone and stress signalling was not always as expected.

For example, 19 SIECs out of 49 which modulated *pDREB2a* activity additionally targeted multiple hormone pathways and we observed a similar number of effectors targeting each hormone pathway that altered abiotic stress or immunity marker expression (Supplementary Figure 1B). The analyses did not reveal a clear correlation between specific hormone and stress signalling functions of SIECs, but rather suggested broader interdependencies with hormone signalling pathways. This might reflect the involvement of hormones in very diverse aspects of plant stress responses (Fahad et al. 2015; Khan et al. 2020; Ku et al. 2018) and the functions of proteins in multiple hormone pathways, of which many have not been described yet (Altmann et al. 2020). While these protoplast assays assigned putative pathway-modulating functions to SIECs, it raised the question of how these proteins might alter these pathways to transduce phenotypic traits in plants.

### Symbiont effectors interact with stress-associated plant proteins

We previously employed yeast-two hybrid analyses to map the SIEC interactome. We identified 33 SIECs interacting with 156 Arabidopsis proteins in the initial yeast-two-hybrid screen (Osborne et al. 2023). To support the biological validity of the SIEC-Arabidopsis interactome, we localized all 33 effector candidates except SIEC54 in *N. benthamiana* and evaluated how frequently SIECs appeared in the same cellular compartment as their 156 target proteins. Confocal microscopy revealed distinct localisations for SIECs *in planta*, including the nucleus, cytoplasm and endoplasmic reticulum (Figure 2A, Supplementary Figure 2).

**Figure 2.**
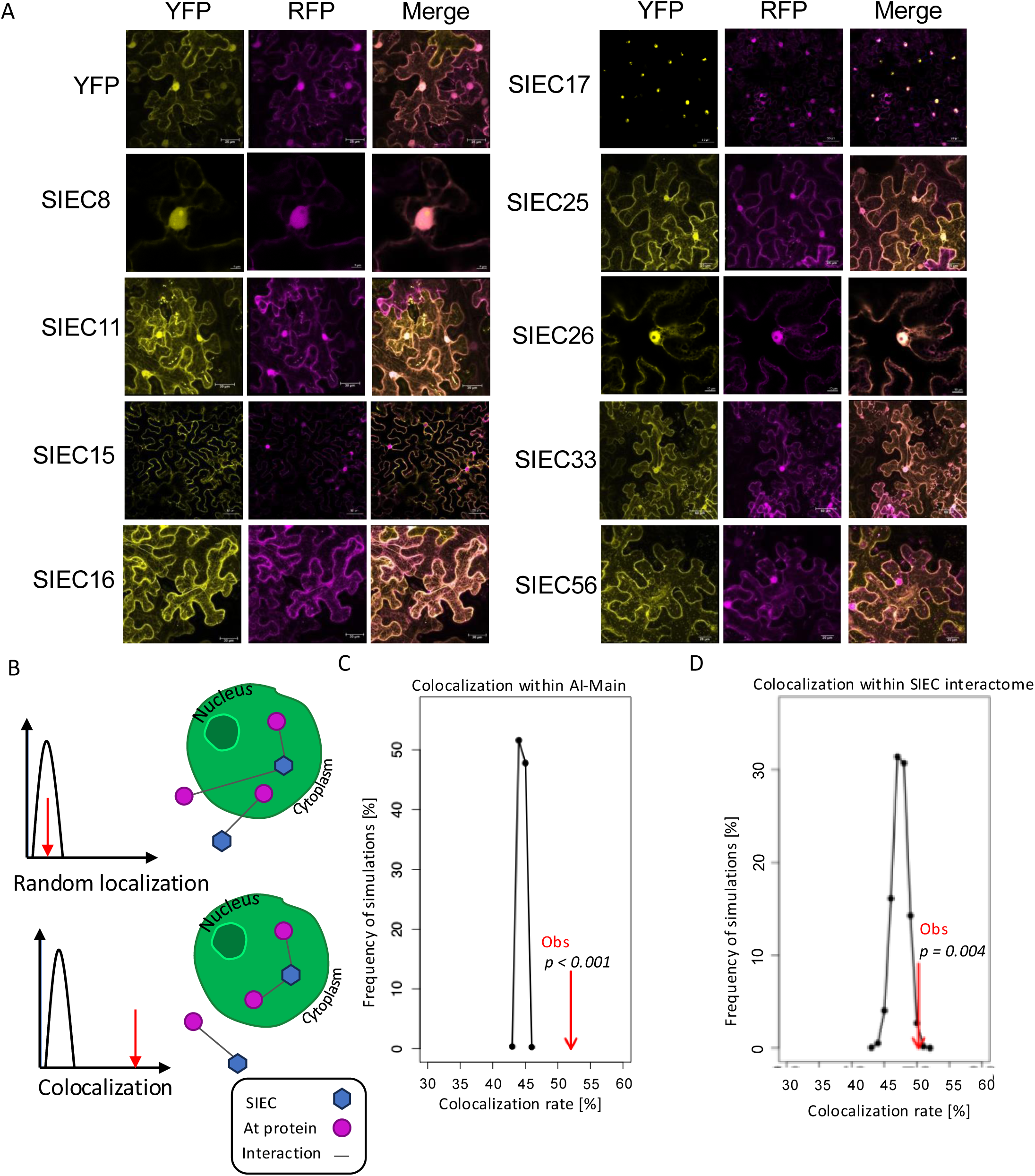
*S. indica* effectors possess distinct subcellular localisations and co-localise with their target proteins *in silico*. **A** Confocal laser scanning microscopy images of SIEC-YFP fusions (yellow) transiently expressed in *N. benthamiana* epidermal cells. Subcellular markers fused to RFP were co-expressed alongside each effector to denote specific organelles (magenta) **B** Network rewiring between an effector (hexagon) and its interacting host proteins (circles) to determine the likelihood of random occurrence in the same cellular compartment. **C** Distribution of the simulated rate of co-localisation in random interaction networks (black points) vs the observed rate in the Arabidopsis Interactome when considering cellular compartment gene ontologies. **D** Distribution of the simulated rate of co-localisation in random interaction networks (black points) vs the observed rate between SIECs (according to CLSM data) and their Arabidopsis targets (GO CC data).

Next, we utilized a network biology approach to predict the likelihood that an effector would appear in the same subcellular compartment as its interacting protein. GO cellular compartment data was first mapped to the Arabidopsis Interactome (AIMain). Under the hypothesis that two interacting host proteins are likely to appear in the same subcellular compartment, we evaluated every pairwise comparison between interacting proteins and their respective localizations, to estimate a colocalization rate of 52% within AIMain. We then randomized the interactions within the network and repeated the analysis, with 10,000 simulated networks. In every iteration, we never observed an equivalent level of colocalization, suggesting our analysis was able to calculate the rate at which proteins within a network colocalize. We repeated this analysis with the SIEC-Arabidopsis interactome data by combining the experimental localization data from *N. benthamiana*, to reveal an estimated colocalization rate of ∼50% (Figure 2 B-D).

Comparative interactomics between SIEC target proteins, and those of pathogen effectors previously revealed that symbiont effectors more significantly interact with the host hormone network (Osborne et al. 2023). To further understand this observation, we investigated the structural and sequence homology of pathogen vs. symbiont effectors to identify specific domains which might influence this activity. We first performed reciprocal protein-BLAST analysis on the protein sequences of effector candidates from *S. indica*, *Hyaloperonospora arabidopsidis* (*Hpa*), *Golovinomyces orontii* (*Gor*), and *Pseudomonas syringae* (*Psy*). Direct statistical comparison of bitscores across all respective alignments revealed no significant differences in homology between effectors of any organism vs. any other (Supplementary Figure 3A), suggesting that effectors are evolutionarily diverse with highly specific functions in line with their respective host ranges. In context of plant-microbe interactions, this suggest an absence of broad-scale homology between effector repertoires of different microbes, aligning with similar observations that have attributed similarity to evolutionary distance (Mukhtar et al. 2011; Weßling et al. 2014). Interestingly however, a PCA of the alignment metadata revealed a symbiont specific shift (Supplementary Figure 3B). Whether the distinction in SIEC sequence homology can be attributed to its broader host range, or its role as a symbiont is unclear. To investigate further, we performed domain analyses of all effectors and evaluated domains which were common and those which were unique between the different organisms (Supplementary Figure 3C, D). SIECs overwhelmingly possessed unique domains which were not present in the pathogen set, including alpha/beta hydrolases, metalloproteases and serine proteases (Supplementary Table 3). We also found that effectors of all organisms possessed intrinsically disorder domains (IDDs). Intra- and interspecies convergence of effectors on host proteins has been previously reported, wherein multiple secreted proteins target the same host proteins, usually to modulate major regulatory hubs (Mukhtar et al. 2011). This raised the question of whether IDDs facilitate convergence by allowing effectors to interact with multiple host targets. We mapped the interactions between *Hpa*, *Gor* and *Si* effectors and Arabidopsis, and classified these based on the presence of an IDD. Analysis of this network suggested IDDs do not facilitate convergence (Supplementary Figure 3E). We observed no statistically significant differences in degree number of effectors based on whether they possessed an IDD, both within and across species. Reciprocal analysis of convergent targets further revealed no link between degree and presence of an IDD in Arabidopsis proteins. Due to the lack of any clear structural distinctions between symbiont and pathogen effectors, we next focused on the function of proteins that interacted with SIECs.

We first curated a database of all Gene Ontologies (GO) for genes which we previously described as “Y2H amenable” across the 8K and 12K spaces, and filtered on terms associated with either biotic, abiotic or general (centering on ROS signalling) stress. These stress annotations were then mapped to the Arabidopsis interactome (AIMain; Arabidopsis Interactome Mapping Consortium 2011, Figure 3A, B). Corroborating previous reports for genes involved in biotic stress, K-shell decomposition analysis of the annotated AIMain network revealed that stress associated genes exist more centrally within the network and represent hubs that are integral to network structure and are biologically important (Figure 3C, Supplementary Figure 4D). Next, we quantified the rate at which microbial effectors (*Si*, *Hpa*, *Psy*, *Gor*) target these stress associated genes. 30, 42, 35 and 15 stress genes were targeted, representing 28%, 34%, 57% and 33% of respective host targets. Degree preserved network rewiring (10,000 simulations) suggested that in each case the interaction between effectors and stress targets was not random, and likely a biologically relevant observation (Supplementary Figure 4A). Given that GO annotations had been classified as biotic or abiotic, we next asked how central stress-associated proteins are within AIMain, and whether different microbes targeted specific groups. Unsurprisingly, stress targets possess a higher K-shell value vs background annotated genes (Figure 4F). This was especially true for convergent targets which interact with effectors from multiple microbes, although previous annotation of these proteins is likely skewed from our previous defining of these genes as functioning in biotic stress responses (Weßling et al. 2014). Remarkably, however, while all organisms interacted with proteins in both categories, we noted that SIECs interacted with more abiotic proteins than pathogen effectors (Figure 3E, Supplementary Figure 4B).

**Figure 3.**
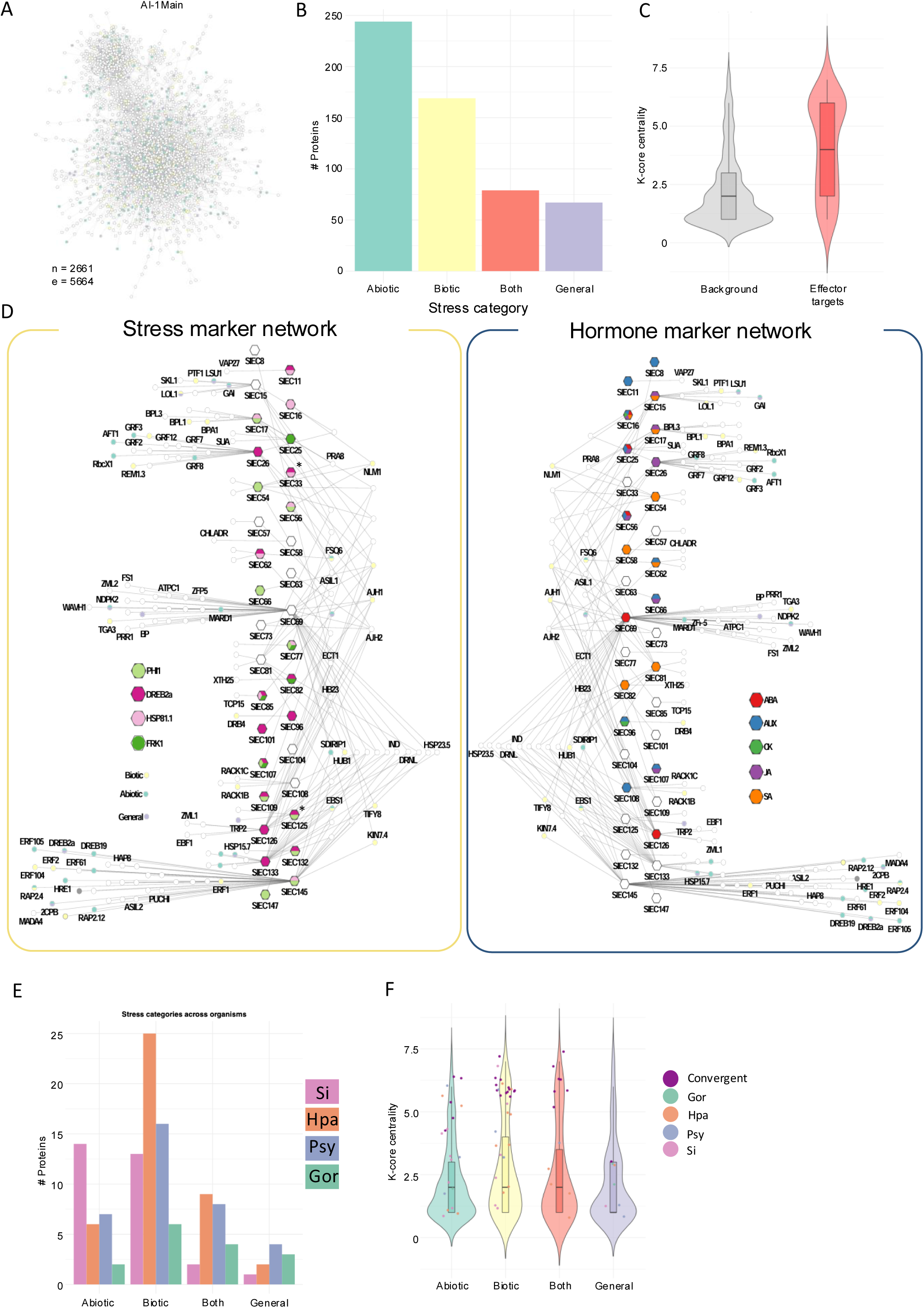
*S. indica* effectors target host stress-responsive proteins. A Network graph of the Arabidopsis interactome (AI-1Main). Proteins with GO terms associated with stress responses are annotated in green (abiotic stress), yellow (biotic stress) or blue (general stress). B Bar plot summarising the frequency of stress annotated proteins within AI-1Main. C Violin plot describing the K-shell distribution of stress annotated proteins within AI-1Main (grey) vs proteins targeted by effectors of *Si*, *Hpa*, *Psy* and *Gor*. D Network graph describing the SIEC-Arabidopsis interaction network. SIECs (hexagons) are coloured according to their function in stress marker assays (A, left) or hormone marker assays (B, right). Interacting Arabidopsis proteins (circles) are coloured according to their known gene ontologies in stress responses. E Bar plot summarising the GO stress annotations of effector targets from *Si*, *Hpa*, *Psy* and *Gor* irrespective of search space. F Violin plot describing the K-shell distribution of stress annotated proteins within AI-1Main based on their functional characterisation as abiotic, biotic, both or general. Individual points represent unique or convergent effector targets from *Si*, *Hpa*, *Psy* and *Gor*.

This prompted us to map our *pSTRESS::LUC* data onto the interactome to identify putative functional relationships between effectors that affect stress responses and interact with proteins that contribute to those responses. Since hormone signalling also supports stress responses, we further mapped our previously reported *pHORMONE::LUC* (abscisic acid, auxin, cytokinin, jasmonic acid, salicylic acid) data onto the interactome. All GO terms for every target where ‘stress’ appeared as a character string in their list of gene ontologies (biological process (BP)) and combined this with the functional data identified in the *pSTRESS* marker screen (Figure 3D). 21 of the 33 SIECs with mapped interactions targeted 55 proteins with GO annotations in stress responses. These included salt, heat, drought and osmotic stress tolerance, as well as defence responses and regulation of plant innate immunity (Supplementary Table 2).

We also observed an overlap between the *pSTRESS*/*pHORMONE* regulatory activity of SIECs in protoplasts and the known function of their targets. For example, SIEC145 modulated the abiotic stress marker *pHSP81.1::LUC* (Supplementary Table 1) and interacted with host proteins involved in heat and water deprivation responses (Figure 3D). SIEC15, which influenced defence hormone signalling in protoplasts, was found to interact with proteins associated with immunity; PLASTID TRANSCRIPTION FACTOR 1 (*PTF1*); TEOSINTE BRANCHED1, CYCLOIDEA AND PCF TRANSCRIPTION FACTOR 13 (*TCP13*) (Wang et al. 2024), and the cell death/hypersensitive response (HR) regulating protein LDS ONE LIKE 1 (*LOL1*) (Ascencio-Ibáñez et al. 2008). In some instances, an overlap was observed in both the *pSTRESS* and *pHORMONE* screens; SIEC17 for instance showed immunity marker regulation (Supplementary Table 1) and affected markers for SA and JA signalling in protoplasts. This effector interacted with BINDING PARTNER OF ACD11 1 (*BPA1*), BPA-LIKE 1 (*BPL1*), and BPA-LIKE 3 (*BPL3*), which are known for their role in plant immunity (Li et al. 2019). Similar observations were made for other SIEC-target combinations, including SIECs 16, 26, 54, 56, 69, 81, 82, 108, 126 and 133, representing 33% of all effectors with mapped interactions.

Taken together, the interaction and functional data revealed symbiont effectors target stress-related host proteins. While the targeting of these proteins appeared not to be unique to symbiont effectors, SIECs targeted more frequently host proteins involved in abiotic stress responses. It suggested that *S. indica* has evolved SIECs to target to alter host abiotic stress signalling. Based on these analyses, we investigated the capacity for six SIECs to confer abiotic stress tolerance in plants, categorized into three groups: SIECs with multiple targets linked to different stress responses (group i), SIECs with a single stress-related target (group ii), and SIECs that modulated *pSTRESS* activity in protoplasts without known targets (group iii) (Supplementary Figure 4E).

### Symbiont effectors modulate abiotic stress response

To analyse SIEC-induced phenotypic changes, we constitutively expressed them under the *35S* promoter in Arabidopsis seedlings. Expression was confirmed by qRT-PCR using SIEC-specific primers and compared to *35S::GFP* expressing control seedlings (Supplementary Figure 5, Supplementary Table 6). We observed enhanced primary root and hypocotyl length in all 6 and in 4 tested *35S::SIEC* lines, respectively (Figure 4A, Supplementary Table 5). Additionally, all (except for *35S::SIEC91*) seedlings produced more lateral roots than *GFP*-expressing control plants (Figure 4C, Supplementary Table 2). As this might be attributed to faster root development, we calculated the average distance between lateral roots. This distance was reduced in *35S::SIEC58* and *35S::SIEC04* plants (Figure 4D, Supplementary Table 5), indicating that both effectors altered lateral root density independent of accelerated root development. Taken together, this analysis revealed that SIECs which regulate stress responses in protoplasts, or target abiotic stress related host proteins, also appear to promote growth in Arabidopsis seedlings.

**Figure 4.**
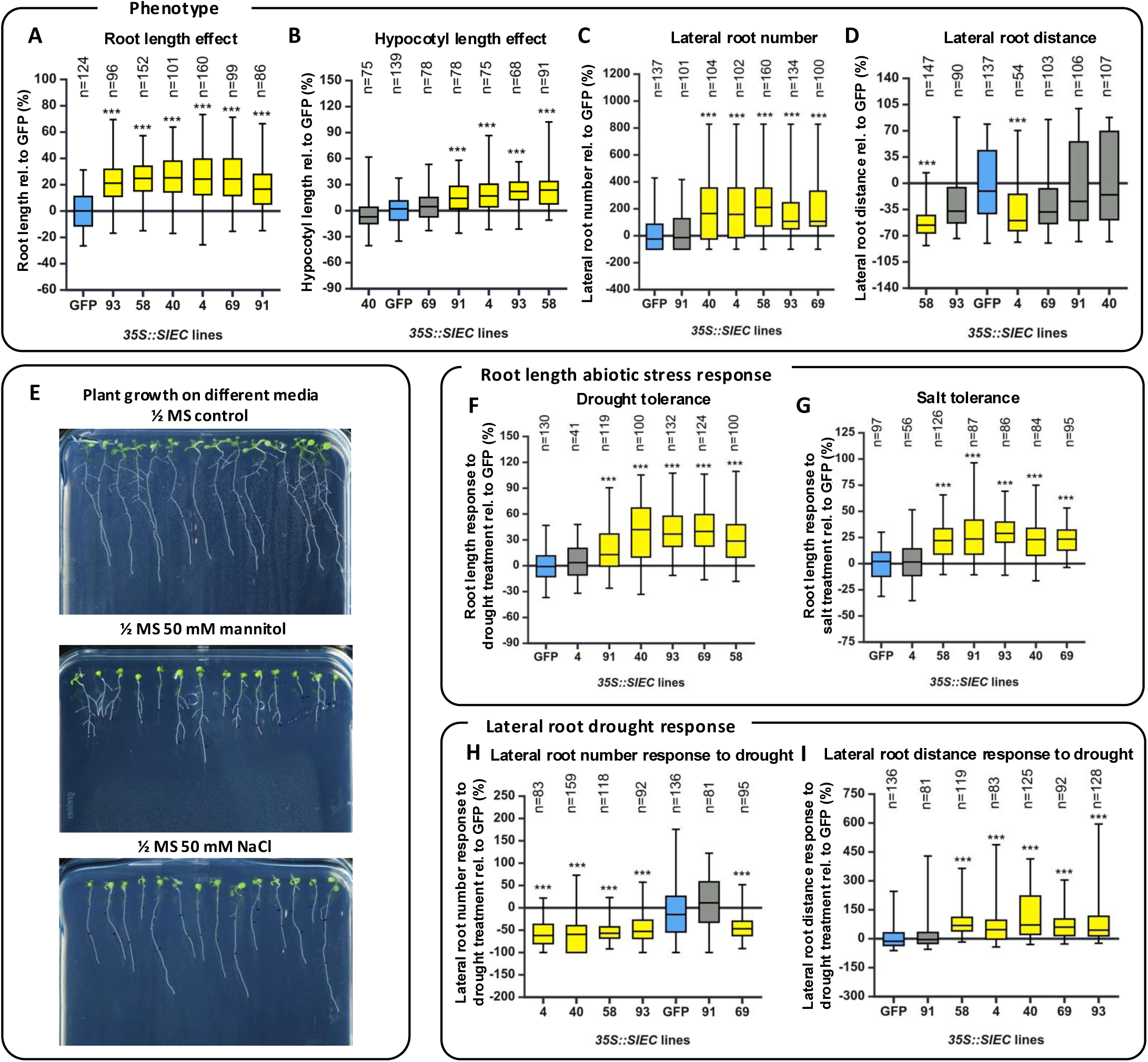
Drought and salt stress phenotyping of *SIEC*-overexpressing seedlings. Roots of 14-day old seedlings treated with mannitol, NaCl or control conditions were measured. Root responses are shown relative to *GFP*-expressing plants (in %). **A-C** Effect of *SIEC* expression on root length (**A**), on hypocotyl length (**B**), on lateral root number (**C**), and average distance between lateral roots (**D**) in control conditions. **E** Representative pictures of plant growth on different stress treatment media. **F-G** Effect of mannitol (**F**) and NaCl (**G**) on root length of *SIEC*-overexpressing seedlings relative to *GFP*-expressing control plants (in %). **H-I** Lateral root phenotypes are altered in *SIEC*-overexpressing plants. Number of lateral roots was counted in mannitol and mock-treated plants (**H**). Lateral root distances were calculated by dividing lateral root numbers by root length (**I**). Both graphs show relative root trait changes in response to mannitol treatment compared to *GFP*-expressing control plants (in %). Positive values indicate increases of root traits as compared to GFP plants. *GFP*-expressing controls are shown in blue; significant effects are indicated in yellow. (*) p < 0.05; (**) p < 0.01; (***) p < 0.001; statistical significance was calculated using a two-tailed, unpaired t-test of at least three replicates. Sample sizes (n) are indicated above each box.

After optimizing experimental conditions (Supplementary Figure 6A, B, Supplementary Table 5), we next monitored root development under salt (NaCl) and osmotic (mannitol) stress. NaCl in plant growth media induces mild salt stress response in Arabidopsis seedling causing reduction of root growth by restraining metabolic processes (Hongqiao et al. 2021). Mannitol leads to osmotic stress similar to drought in plants by decreasing the osmotic potential of the media resulting in decreased primary root length and lateral root development (Lipavsk and Vreugdenhil 1996) (Figure 4E).

By calculating the relative primary root length reduction between control and stress conditions of *35S::SIEC* and *35S::GFP* plants, we found that 5 of the 6 effectors improved root length under mannitol-induced osmotic stress (SIECs 40, 58, 69, 91 and 93; Figure 4F, Supplementary Table 5). Conversely, the relative rate of lateral root induction was suppressed in *35S::SIEC* lines (SIECs 4, 40, 58, 93, 69) as compared to control seedlings, although this might be due to the existing increased rate of lateral root formation in these genotypes (Figure 4 H, I). Similarly, primary root growth was less impaired in the same genotypes under salt stress. (Figure 4G, Supplementary Table 5). Interestingly, SIEC04 did not alter primary root length under drought or salt stress (Figure 4F, G), despite increasing root length and hypocotyl growth under control conditions (Figure 4A, B). In sum, these root phenotyping analyses revealed effector candidates altered plant root length under salt and simulated drought stress thereby confirming our predictions based on our protoplast marker assay and network analysis.

To assess whether the protective effect of SIECs against abiotic stress is related to their interacting proteins, we also analyzed the root responses of respective knockout plants to salt and drought stress. For example, SIEC69 interacts with ZFP5 and TCP9 (Osborne et al. 2023), and promoted primary root length under control and abiotic stress conditions (Figure 4A,F-G) when expressed in Arabidopsis. Conversely, *tcp9* and *zfp5* mutants produced shorter roots and also exhibited altered responses to salt treatment (Supplementary Figure 7). These results indicate that the observed beneficial growth and stress response effects of *SIEC69* expression in Arabidopsis might be related to the function of its targets. Furthermore, this analysis has revealed putative functions for ZFP5 and TCP9 in regulating root growth and abiotic stress responses. Other candidates identified from the interactome include DRB4 (SIEC96 target) and ASIL1 (SIEC107 target). While SIEC107 modulated markers for both *pSTRESS* and *pHORMONE* (Figure 3D), the stress responses of *asil1* plants were equivalent to control plants. Nonetheless, root length was reduced in this mutant (Supplementary Figure 7). Finally, *drb4* knockout resulted in shorter roots, and improved root length response to both salt and drought stress (Supplementary Figure 7); connecting our observation that *SIEC96* enhanced *pDREB2a::LUC* expression (Supplementary Table 1). Overall, by analyzing the growth and stress responses of selected effector target knockout plants, we confirmed that the functions of SIECs *in planta* appear to align with those of their interacting plant proteins.

### Symbiont effectors modulate plant pathogen resistance

Given that some effectors were able to modulate transcriptional PTI responses (*pFRK1, pPHI1)* and defence hormone signalling (*pWRKY70*, *pJAZ10*), we next tested whether SIECs altered seedling susceptibility to different phytopathogens. Unlike the abiotic stress screen, which was directed by the network analysis, we took an undirected approach for the biotic stress screen. Available *SIEC* expressing Arabidopsis genotypes (95 *SIEC* expressing Arabidopsis lines were available for pathogen infection assays, 86, 50 and 70 were tested for *Phytophthora parasitica*, *Botrytis cinerea* and *Rhizoctonia solani* infection, respectively) were screened for altered resistance against three pathogens with different life strategies and target tissues. In addition to the leaf and root infecting oomycete *Phytophthora parasitica,* the leaf infecting fungus *Botrytis cinerea* and the root infecting fungus *Rhizoctonia solani*. While both fungi have a necrotrophic life style, the oomycete is a hemi-biotroph that initially colonizes living cells before switching to necrotrophy (Meng et al. 2014; Choquer et al. 2007; Ajayi-Oyetunde and Bradley 2018).

10 lines possessed altered resistance phenotypes against *P. parasitica* with 5 showing increased (SIEC 30, 33, 47, 96 and 99) and 5 reduced infection (SIEC24, 90, 119, 125 and 130, Figure 5A, Supplementary Table 5, 8). We identified *SIEC08* and *SIEC19* expressing plants as resistant to *B. cinerea* infection, while *SIEC21* expressing plants were more susceptible (Figure 5B, Supplementary Table 5, 8). Only *SIEC82* conferred resistance to *R. solani*, while *SIEC46* increased susceptibility (Figure 5C, Supplementary Table 5, 8). Interestingly, no SIEC altered resistance to more than one pathogen, suggesting their functions are highly specific and might be explained by the target tissue (roots, leaves) and trophic strategy of each pathogen. 87% (13/15) of SIECs which altered pathogen susceptibility in whole plants also altered *pFRK1*, *pPHI1* (PTI markers), *pWRKY70* (SA marker) or *pJAZ10* (JA marker) activity in protoplasts, demonstrating the predictive power of this assay (Supplementary Table 1). 4 SIECs also had targets with functions related to stress responses; SIEC08 interacted with the defence related protein KINESIN 7.4 (KIN7.4) (Shimono et al. 2016), SIEC82 with the defence and SA response associated protein XYLOGLUCAN ENDOTRANSGLUCOSYLASE/HYDROLASE 25 (XTH25) (Ascencio-Ibáñez et al. 2008; Li et al. 2004), SIEC96 with the dsRNA binding DOUBLE-STRANDED-RNA-BINDING PROTEIN 4 (DRB4), potentially involved in viral defence (Qu et al. 2008; Jakubiec et al. 2012), and SIEC125 with the oxidative stress related protein TRYPTOPHAN BIOSYNTHESIS 2 (TRP2) (Zhao et al. 1998). In addition to the abiotic stress phenotyping, these infection assays revealed that SIECs have diverse functions in stress adaptation in plants.

**Figure 5.**
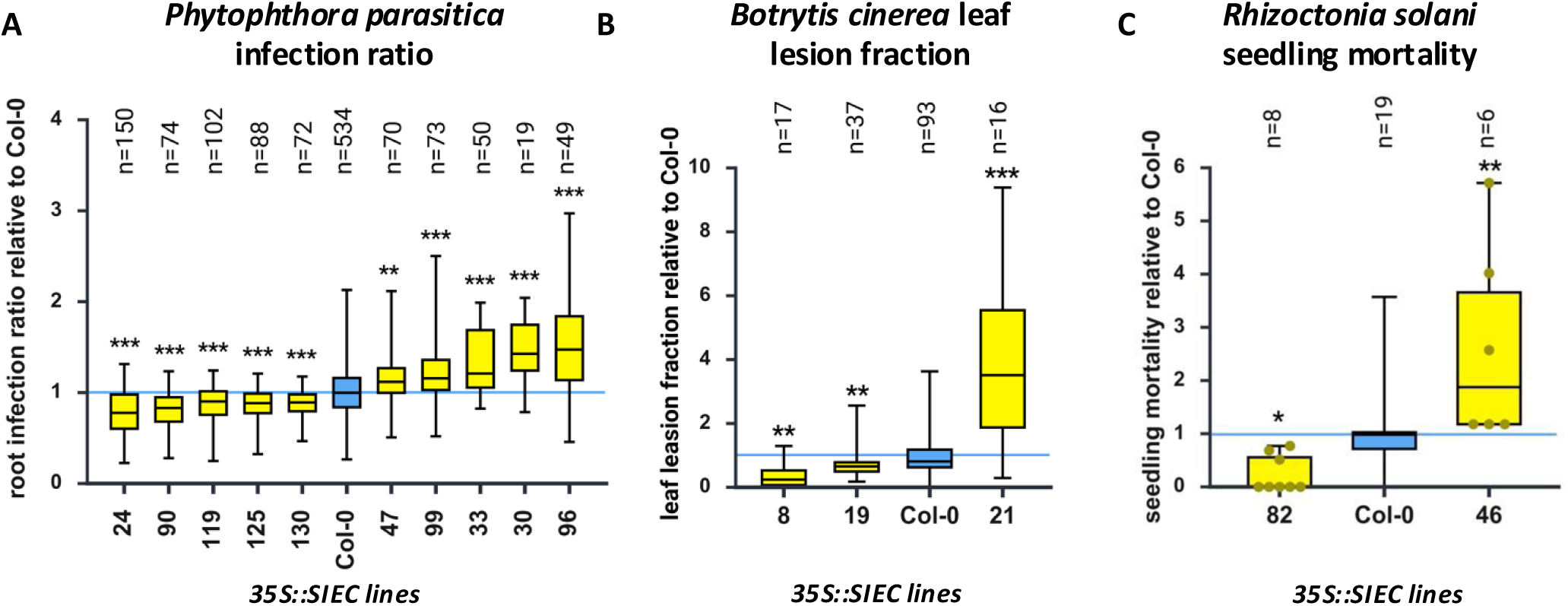
Pathogen infection response of *SIEC* expressing plants. **A-C** *SIEC*-expressing plants were analysed after infection with pathogens. Only significant results are shown in this figure relative to Col-0 infection for *Phytophthora parasitica* root infection ratios (**A**), *Botrytis cinerea* leaf lesion fraction (lesion area relative to the whole leaf area) (**B**), and *Rhizoctonia solani* seedling mortality following root infection (**C**). Col-0 control plants are indicated in blue; significant effects are indicated in yellow. (*) p < 0.05; (**) p < 0.01; (***) p < 0.001; statistical significance was evaluated by two-tailed, unpaired t-test of at least two biological replicates. Sample sizes (n) are indicated above each box.

## Discussion

It has long been understood that phytopathogens utilize secreted proteins to suppress host immunity and alter host signalling and metabolism to support their infection and reproduction (Hogenhout et al. 2009). These interactions are generally deleterious to host fitness and negatively impact host responses to abiotic stresses, growth and yield. In addition to manipulating host signalling, recent studies revealed that beneficial and pathogenic plant symbionts release effectors to directly manipulate plant-associated microbiota. The antimicrobial activity of these “ecological” effectors can shape the composition and thus the effect of microbial communities on plants (Snelders et al. 2022; Mesny et al. 2026). Interestingly, *Si* and close relatives can activate beneficial effects in host plants in the absence of any other microbe or microbial community (Waller et al. 2005; Deshmukh et al. 2006). Based on this rationale, we hypothesized that effectors from *Si* might act to enhance, rather than hinder, host fitness.

*Si* increases plant growth under regular soil nutrition, and improves phosphate (Kumar et al. 2011; Yadav et al. 2010), iron (Verma et al. 2022) and sulfate (Narayan et al. 2021) uptake to its host. In addition to altered nutrient supply, *Si* enhances salt stress tolerance, associated with an improved antioxidant status (Baltruschat et al. 2008; Waller et al. 2005), growth under heat stress through the modulation of JA and ethylene signalling (Chen et al. 2022), and biotic stress tolerance through induced systemic resistance, for example, by modulated gene expression (Molitor et al. 2011). Stimulation of antioxidative metabolism and/or stress-related genes might also be involved in increased tolerance to other stresses (Gill et al. 2016), including chilling (Jiang et al. 2021), pathogen defence (Stein et al. 2008) and drought (Zhang et al. 2018). A natural hypothesis to these diverse benefits is that increased nutrient uptake potentiates systemic host fitness against diverse environmental factors. However, the priming of adaptive genes or modulations to antioxidative metabolism in *Si* colonized plants cannot be attributed solely to improved nutrient acquisition. We therefore hypothesized that an essential factor in plant-microbe symbiosis is the utilization of effector proteins to “fine-tune” host signalling which confers plant benefits.

Consistent with this, we recently identified SIEC141 to interact with thioredoxin-like protein CDSP32 resulting in the activation (monomerization) of NON-EXPRESSOR OF PR1 (NPR1), a key regulator of systemic immunity (Zhang et al. 2024). Here, we show that SIECs, target, and in some cases alter response to biotic and abiotic stresses. Screening effector functions in Arabidopsis leaf protoplasts revealed that 82/106 SIECs (77%) altered either basal or inducible transcriptional activity of fluorescent markers for flg22-mediated PTI or heat stress. By analysing our previously reported SIEC-Arabidopsis interactome in more detail, we found that SIECs interact with more host proteins associated with abiotic stress as compared to pathogen effectors and possess unique domain architecture (e.g. alpha/beta hydrolases, metalloproteases, serine proteases; Supplementary table 3) which may represent a symbiont specific fingerprint. To support our interactome, we localized all except one SIEC in *N. benthamiana* and found that their subcellular localisations reflected the localisations of their target proteins (Figure 2A, Supplementary Figure 2). While this approach shows that SIEC possess distinct subcellular localisations, we recognise that transient expression of effectors in N. benthamiana does not reveal the spatiotemporal dynamics of effector secretion during a bone-fide Si-Arabidopsis interaction in the root. Given the technical challenge of genetically engineering *Si* to observe effector secretion and subcellular localization, this method allowed us to evaluate the likelihood that two proteins would at least appear in the same organelle and revealed a higher rate of colocalization between effectors and their targets, than occurs in random networks.

Based on our protoplast and interactome data, SIECs appear to exert overlapping hormone and stress activities, suggesting that their functions in stress signalling are coupled to their hormone-modulating activities (Figure 1G, Supplementary Figure 1). ABA, as a major regulatory hormone in abiotic stress responses, for example, integrates extrinsic signals and redirects plant development to cope with drought, high temperature and salinity (Hong et al. 2013). These functions are often mediated in crosstalk with other hormones, such as auxin, which regulates lateral root branching (Seo and Park 2009). Of the six SIECs tested *in planta*, five improved drought stress response and changed ABA marker activity in protoplasts (Figure 4F, Supplementary Figure 4E). Plants expressing these effectors displayed increased lateral root distance and root length after drought exposure (Figure 4F-I). SIEC04 altered ABA and AUX pathways in protoplast assays (Supplementary Figure 4E) and affected lateral root responses to mannitol. Furthermore, SIEC69 and SIEC58 enhanced root length and reduced lateral root density in response to drought stress as compared to control plants (Figure 4A, F, H-I). Interestingly, expression of *SIEC69* modulated AUX and JA, in addition to ABA markers, and interacted with multiple stress related proteins (Figure 3D, Supplementary Figure 4E), suggesting it may have broad regulatory roles in the *Si*-Arabidopsis symbiosis. This is further reflected in the interactome, where this protein acts as a major hub with 44 putative host interactions (Osborne et al. 2023). SIEC58 affected ABA and SA pathways in plants (Figure 3D) and interacted with CHLOROPLAST ALDEHYDE REDUCTASE (CHLADR, AT1G54870; Supplementary Figure 4E), which is involved in detoxification of reactive carbonyls (Yamauchi et al. 2011). These results indicate that effectors not only increase drought tolerance by promoting root growth but also affect root architecture (e.g. lateral root formation). We further observed that these SIECs reduced the sensitivity of roots to salt stress-mediated growth arrest (Figure 4G). Here, we focused on understanding seedling responses to osmotic and salt stress. While it is also important to distinguish between sensitivity and survival in these assays, our phenotyping data allude to potential functions of specific effectors in promoting fitness under abiotic stress. Moreover, our data suggest that SIECs and their respective targets converge onto similar physiological and developmental processes to regulate growth and stress responses (Figure 3D). Given that both SIEC107 and ASIL1 affect auxin responses (Osborne et al. 2023; Gao et al. 2009) and are nuclear localized (Supplementary Figure 2, Supplementary Table 9), suggests effects on auxin-dependent transcriptional programs that regulate growth, rather than stress responses (Supplementary Figure 2). The observed beneficial functions of *Si* effectors might depend on the activities of their respective targets and strengthens our rationale that investigating the activity of symbiont effectors can reveal how existing host pathways are fine-tuned to mediate both mutualism and systemic benefit. Combined with our previous study, this also highlights the suitability of using SIECs to detect previously unknown crosstalk points between stress and hormone signalling networks that are critical for plant stress resilience.

In addition to altering abiotic stress response, *Si* increases host resistance to pathogens (Stein et al. 2008; Sun et al. 2014; Waller et al. 2005). Several SIECs were shown to modulate host immunity in both protoplasts and whole seedlings. Among the SIECs affecting pathogen resistance, the majority altered root colonization by *P. parasitica*. This might reflect similarities in the infection strategies and the target tissue (roots) of *P. parasitica* and *Si* (Jacobs et al. 2011; Meng et al. 2014). Significant effects for *R*. *solani* infection were observed less frequently, possibly due to the small sample size and the harsh conditions of seedling mortality assays, making it difficult to detect moderate changes. Nonetheless, we identified eight SIECs that decreased, and seven that increased disease susceptibility (Figure 5A-C, Supplementary Table 8). While this number is lower than the number of effectors which modulated PTI signalling in protoplasts, we believe this might suggest tissue specific activity of some effectors in roots vs shoots, or a potential limitation to screening fungal effectors for altered responses to flg22-mediated activation of immunity. Interestingly the effectors which altered susceptibility possessed functions in hormone and/or stress signalling in protoplasts or were found to interact with stress related Arabidopsis proteins (Figure 3D, Supplementary Table 1,2). This included altered expression of markers for SA and JA as major plant defence regulating hormones (Hou and Tsuda 2022), in addition to auxin and cytokinin with key functions in growth and development (Brumos et al. 2018; Li et al. 2021). While the latter hormones are not typically linked to pathogen defence, considerable crosstalk between hormone pathways connect auxin and cytokinin to plant immunity, especially in balancing growth and defence (Naseem et al. 2015). It is therefore unsurprising that SIECs which modulate hormone signalling also affect PTI signalling and pathogen susceptibility.

Both pathogens and symbionts activate host immunity, which must be overcome for successful colonization (Jacobs et al. 2011; Yu et al. 2019; Enebe and Erasmus 2023). *Si* suppresses host immunity and might use SIECs for this purpose. Consistent with this, PIIN_00029 was previously identified as an *Si* effector-like small-secreted protein with sequence homology to E3 ligases. It was suggested that this protein increases host colonization by suppressing plant defence and ROS generation (Plett and Martin 2015; Zhang 2014). In addition, *Si* effector PIIN_08944 was reported to interfere with SA homeostasis to suppress PTI, SA-related defence gene expression and ROS burst (Akum et al. 2015). *Si* effector Dld1 (PIIN_05872) potentially enhances micronutrient accessibility for the microbe and interferes with ROS homeostasis to aid colonization (Nostadt et al. 2020). While these effectors were not present in our top 106 candidates, they illustrate the necessity for *Si* to suppress host immunity. This is perhaps counterintuitive given that *Si* colonization can induce systemic resistance within 19 days of colonization (Molitor et al. 2011), and that some SIECs confer disease resistance (Figure 5A-C). It is, however, explainable in the context of temporal interaction dynamics. *Si* releases SIECs in a stage-dependent manner, with early effectors delivered during the onset of interaction to inhibit immunity, and late effectors to enhance resistance upon successful accommodation. Characterizing these secreted proteins will be essential to understand their roles in facilitating either colonization, mutualism, or both; especially during the *bona fide Si*-host interaction.

Just as pathogen effectors have been used to decode plant immunity, we hypothesize that symbiont effectors can be studied to understand symbiosis and identify beneficial host pathways that promote fitness. Here, we not only revealed SIECs with immune suppressing functions, but also those which modulate plant responses to abiotic stress. They are therefore a valuable resource to establish regulatory principles and mechanisms of plant beneficial traits within complex, often crosstalking plant signalling networks. For some *Si* effectors we could neither identify a host target nor detect any effects in our plant signalling assays. It will be interesting to further our understanding to what extent *Si* effectors exert unique or additional antimicrobial activities as recently reported for plant pathogen effectors (Mesny et al. 2026). Given its broad host range, we hypothesize that *Si* effectors offer a sustainable strategy for identifying breeding targets of beneficial signalling pathways in crop species, particularly given the impact of climate change on agricultural productivity (Fischer et al. 2021; Lobell and Gourdji 2012).

## Materials and Methods

### Experimental model and subject details

All *Arabidopsis thaliana* (Arabidopsis) mutants and transgenic lines are in the Col-0 accession (wild type, WT). *A. thaliana* genetic material including *35S::SIEC* and t-DNA insertion target knockout lines were described previously (Osborne et al. 2023).

Growth conditions for specific experiments are given below

### Protoplast screening

Protoplast screening was performed as described in Osborne et al. 2023, following the procedure in Lehmann et al. 2020. 4-5-week-old Col-0 Arabidopsis plants were used for mesophyll protoplast generation. Following these protocols 10,000 protoplasts were transformed with 3 µg DNA. Each transformation contained 3 different plasmids: 1 µg *pUBQ10::GUS* (normalization control), 1 µg *pSTRESS::LUC* (luminescent marker for stress pathway analysis) and 1 µg *p35S::SIEC* (*SIEC* expression). Empty vector expressing Nt-HAEMAGLUTATIN instead of *SIEC* expressing ones were used as control. After transformation and over-night incubation protoplasts were treated according to the transformed stress marker. For abiotic stress markers *pDREB2a::LUC* or *pHSP81.1::LUC* transformations heat stress was applied for 1 h at 37 °C or kept in short day conditions as basal control. Protoplast transformed with PTI markers *pPHI1::LUC* or *pFRK1::LUC* were treated with 0.1 mM flg22 or water as basal control. To indicate SIECs with function in signalling pathways mock and treatment ratios were multiplied to show effects in the same direction and reveal promising signalling modulating candidates.

### Interactome and stress-signalling network analysis

The SIEC-Arabidopsis interactome was previously described (Osborne et al. 2023). To identify host targets associated with stress, we mapped all biological process gene ontologies for all proteins within the Arabidopsis Interactome (AI1Main) (accessed 21.08.25). Ontologies were then subcategorized into their association with biotic, abiotic or general stress responses based on the ontology itself, or its parent GO term. General stress responses comprised of terms associated with oxidative stress, DNA damage repair, the ERAD pathway, and reactive oxygen species metabolism, as these terms - while involved in plant stress responses - cannot be described as exclusively biotic or abiotic. Classification of proteins within the interactome allowed the comparison between effector targets of *Si*, *Hpa*, *Gor* and *Psy*. To calculate the random likelihood that any organisms’ effector repertoire would target stress proteins at random, degree preserving network rewiring was performed. In short, given X effectors target Y host proteins, Y proteins were sampled randomly from all available proteins with interactions established across our Y2H platform (Y2H Ammenable). The number of stress associated proteins was then stored (Z) and this analysis was repeated 10,000 times (N). p-values were calculated based on how many simulated networks had more stress associated proteins than was observed experimentally:

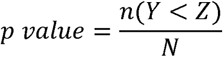

K-shell analysis was performed using the R package “igraph”. Network K-cores were computed using the function “coreness” to evaluate node centrality within AI-Main. Stress annotations were then mapped to respective K-shells to evaluate centrality of those proteins.

### Effector domain and homology analysis

To identify organism specific effector domains, proteins sequences of all effectors from *Si*, *Hpa*, *Gor* and *Psy* were analysed using the EBI InterPro tool (https://www.ebi.ac.uk/interpro/) with default settings. Domains were then compared between effector sets to evaluate occurrence of specific domains between organisms. Statistically significant differences between effectors either with or without intrinsically disordered domains were calculated by Wilcoxon test.

For sequence homology analysis, the NCBI BLASTp tool was used (https://blast.ncbi.nlm.nih.gov/Blast.cgi). Reciprocal BLASTp analysis was performed between effector protein sets, using every combination of query and reference (e.g., *Si* as reference vs. *Hpa* as query, and *vice versa*). A PCA was performed comparing bitscores across all comparisons.

### Gene expression analysis by quantitative real time-PCR

A pool of seedling roots was frozen in liquid nitrogen and ground into a fine powder. Total RNA was extracted using TRIzol^®^ reagent (Invitrogen). RNA was digested by DNase I (Thermo Fisher Scientific) including RiboLock RNase Inhibitor (Thermo Fisher Scientific) to remove genomic DNA. Following manufactures instructions cDNA syntheses was performed using qScript™ cDNA Synthesis Kit (Quantabio). For expression analysis in qRT-PCR SYBR^®^ Green JumpStart™ *Taq* ReadyMix™ (Sigma) was used following standard protocol. To determine absolute and relative expression of *SIECs* the 2^-ΔCt^ and 2^-ΔΔCt^ methods were used (Livak and Schmittgen 2001). Primers used for this analysis are listed in Supplementary Table 7.

### Stress response assays

Arabidopsis seedlings expressing *35S::SIEC* and target knockout t-DNA plants were phenotyped for root growth and lateral roots in control and stress conditions. Stress response was analysed by comparing mock vs treated root length, lateral root number and density (%) to *GFP* expressing or Col-0 control plants. Seeds were initially surface sterilized using NaOCl solution and sown on ½ MS media, containing 10 µg ml^-1^ BASTA for *SIEC* expressing plants, following stratification for 48 h at 4 °C. Selection on BASTA and germination took place for 7 d in short day conditions in growth cabinets at 22 °C. Seedlings in similar developmental stages were then transferred on ½ MS media containing 50 mM mannitol, 50 mM NaCl or no treatment. After another 7 days growth in short day conditions pictures were taken and root length and lateral root number was measured using ImageJ. Stress response was quantified by calculating the percentual change of root length / lateral root number / lateral root distance between mock and treatment conditions. Lateral root distance was calculated by dividing the average root length by lateral root number. These values were compared to the percentual change of *35S::GFP* or Col-0 plants, by dividing *35S::SIE*C or target knockout % change by the mean of the respective control % change to show the percentual difference in treatment response. From these values 100 was subtracted to visualize reduced effects with negative values, while positive values indicate increase.

### Effector localization in *N. benthamiana* epidermal cells

Effector subcellular localization was first predicted using the https://localizer.csiro.au/ tool (Sperschneider et al. 2017), which classifies compartment targeting to the cytoplasm, nucleus, chloroplast or mitochondrion based on primary sequence analysis. Next, *Agrobacterium tumefaciens* mediated transient transformation was used to express respective YFP-SIEC fusions in *N. benthamiana* cells. 2 days after infiltration, leaf discs were imaged using Leica SP8 confocal laser microscope with excitation at 514 nm and detection between 520-545 nm. To compute the random likelihood that effectors and their targets would appear in the same subcellular compartment, gene ontologies for cellular compartment (accessed 02.05.25) were mapped to all effector targets and proteins within the Arabidopsis interactome. To make our analysis more stringent, we only included host proteins which localised within the cytoplasm, nucleus or endoplasmic reticulum. The colocalization rate was calculated by considering all localisations between proteins within respective interaction networks. For proteins with multiple predicted localizations, all pairwise compartment comparisons were assessed. For example, if protein X is localized exclusively to the nucleus and protein Y to both the nucleus and cytoplasm, their overlap in the nucleus corresponds to a 50% localization rate. Degree preserving network rewiring was performed to evaluate the likelihood two proteins would appear in the same compartment at random https://doi.org/10.5281/zenodo.7749043 (Osborne 2023). Interactions within networks were randomised, while protein localisations were maintained.

### Pathogen infection assays

#### Phytophthora parasitica

*35S::SIEC* plants were grown vertically on sterile ½ MS plates containing sucrose in a 13 h light, 11 h dark cycle at 23°C for 10 days. Seedlings were then transferred onto sucrose-free ½ MS plates for *P. parasitica* infection assays. *P. parasitica* transformant Pp1121, which stably expresses GFP was used for inoculation (Wang et al. 2011). Fresh *P. parasitica* mycelial plugs (5 mm in diameter) cut from the edge of a culture grown on carrot agar (5% (v/v) cleared carrot juice agar (CA) medium supplemented with 0.002% (w/v) β-sitosterol and 0.01% (w/v) CaCO_3_) in the dark at 23 °C for 3-5 days were transferred onto the middle of a root of each individual Arabidopsis seedling. The colonization of the *P. parasitica* hyphae along plant roots was observed using a fluorescence microscope at 48 hpi. The length of the infected root part and the total root length were measured using ImageJ.

#### Botrytis cinerea

*B. cinerea* (strain Bc001) was isolated from tomato and cultured on potato dextrose agar (PDA) in a growth chamber at 16 °C for 15 days. Spores were harvested from mycelia with 3 ml potato dextrose broth (PDB) and subsequently were filtered through four layers of gauze to remove hyphae. Spore density was adjusted to 5 × 10^5^ spores per ml with PDB. Fully expanded leaves of 4-week-old Arabidopsis seedlings were detached and infected by dropping 5 μl of the spore suspension. Lesions were measured at 48 hpi.

#### Rhizoctonia solani

*R. solani* (strain HBZJ-5X) was cultured on PDA in a growth chamber at 23 °C for 7 days. 10-day-old Arabidopsis seedlings vertically growth on ½ MS (with sucrose) media were transferred onto sucrose-free ½ MS media for subsequent inoculation. Mycelium plugs (diameter 5 mm) were harvested from *R. solani* PDA culture and put on the middle of individual Arabidopsis seedlings roots. The inoculated seedlings were vertically grown at 23 ℃ with an 11/13 h day/night photoperiod for 9 days.

All graphs involving biotic and abiotic stress phenotyping were created in https://BioRender.com.

## Acknowledgements

This work was funded by research grants from Biotechnological and Biological Research Council (BBSRC) / Engineering and Physical Sciences Research Council Grant (EPSRC) of the United Kingdom (BB/M017982/1) and BASF Plant Science Company GmbH.

## Author contributions

Research concept and design: L.R., R.O., S.L., P.F.-B., W.S. and P.S. SIEC-At interactome mapping and data integration: R.O. Protoplast assays: L.R., R.O., S.L., J.R., and E.K. Generation of SIEC lines and stress phenotyping: L.R., R.O., S.L., E.K., J.R., R.E., and Y.Z. Manuscript writing: L.R., R.O., L.R., S.L., P.F.-B., W.S., and P.S.

---

**Supplementary Figure 1.**
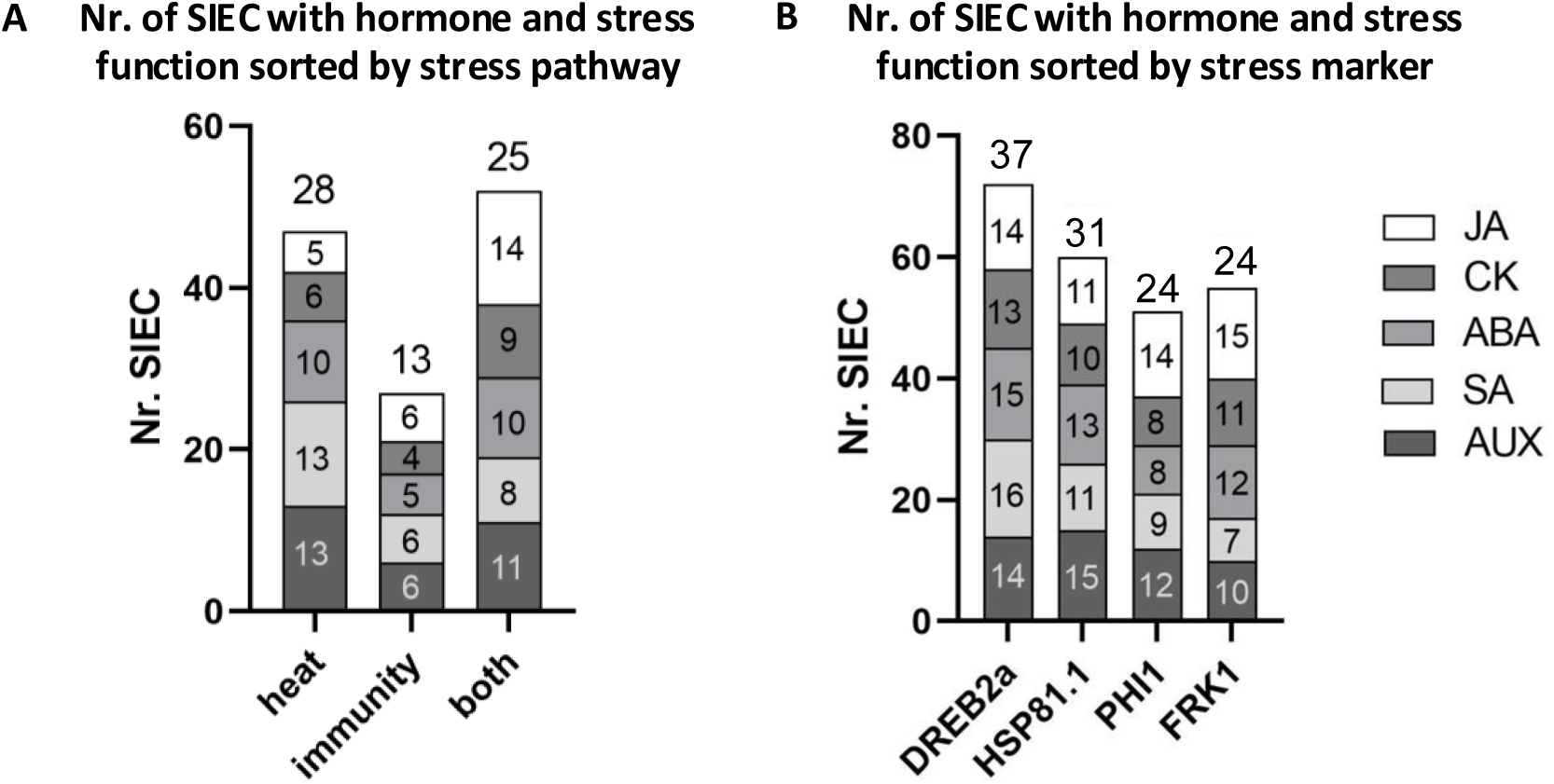
Number of SIECs with stress and hormone function. **A** Number of SIECs that alter stress marker expression and have additional hormone functions. **B** Number of SIEC that specifically modulate stress signalling and, in addition, hormone pathways. The number above each bar indicates the total number of SIECs which is lower than the sum as individual SIECs can affect multiple hormone and stress pathways. AUX – auxin, CK – cytokinin, ABA – abscisic acid, JA – jasmonic acid, SA – salicylic acid

**Supplementary Figure 2.**
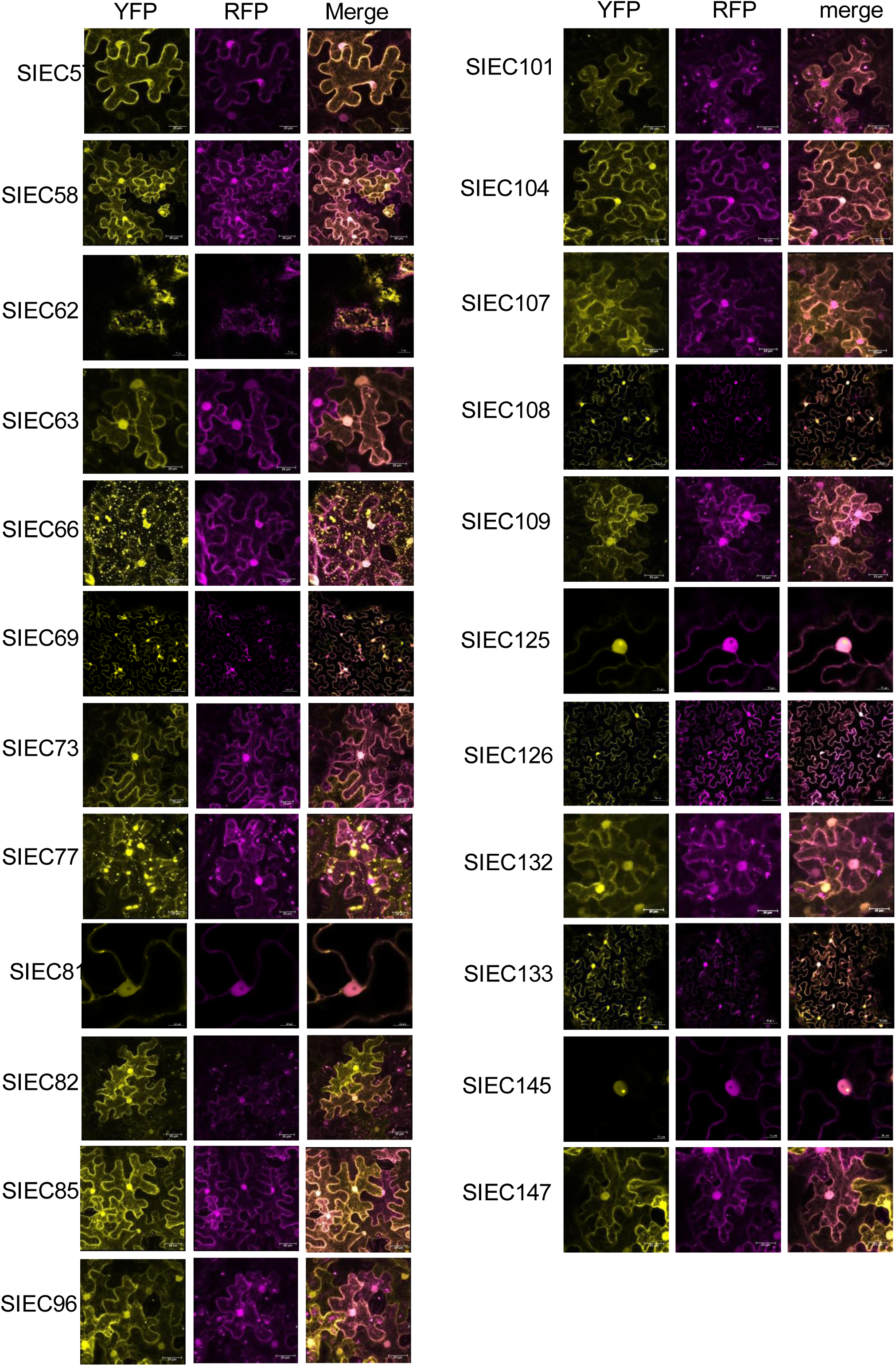
Confocal laser scanning microscopy images. Confocal laser scanning microscopy images of SIEC-YFP fusions (yellow) transiently expressed in *N. benthamiana* epidermal cells. Subcellular markers fused to RFP were co-expressed alongside each effector to denote specific organelles (magenta).

**Supplemental figure 3.**
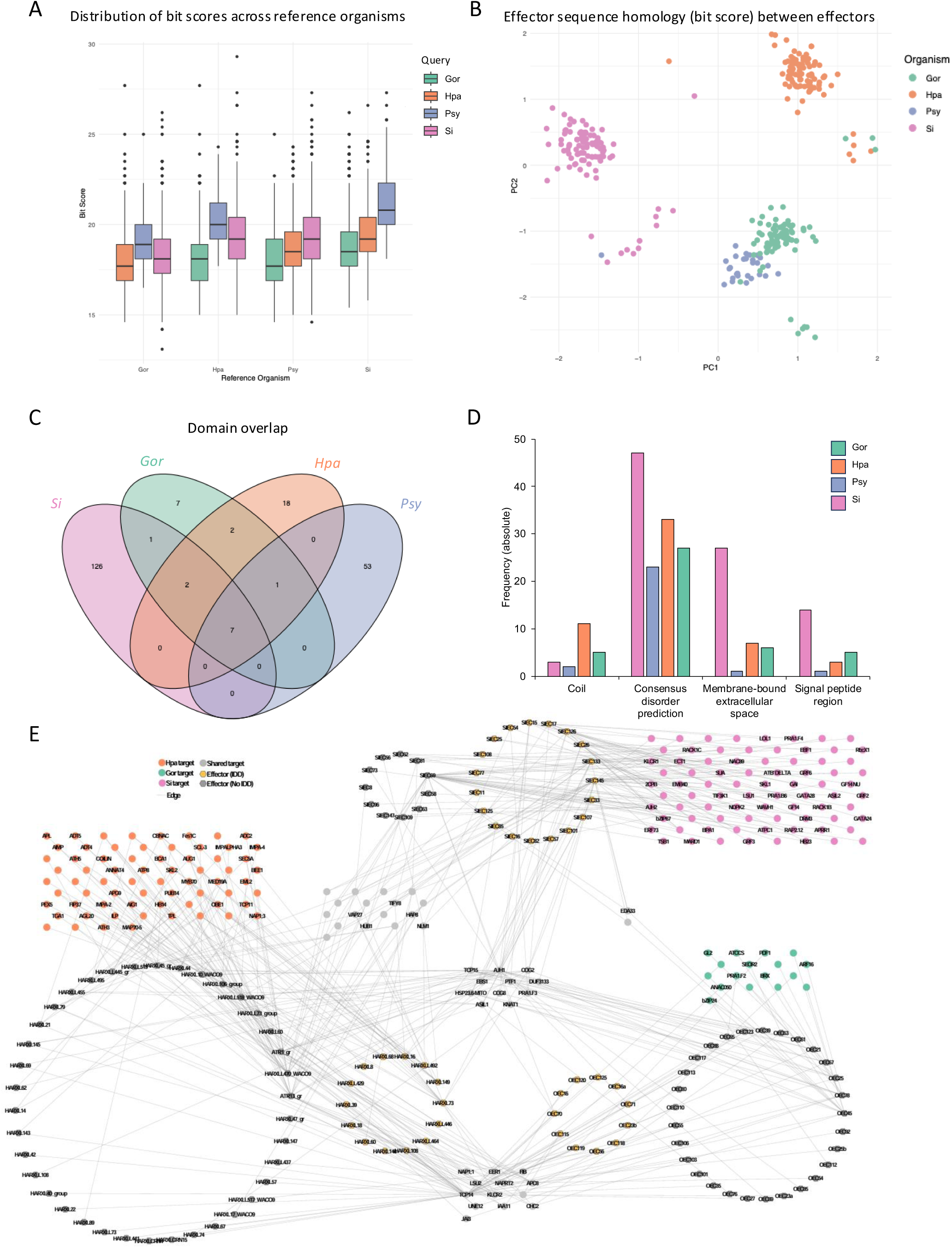
Sequence and protein domain analysis of microbial effectors. **A** Box and whisker plots showing Bit score across reciprocal pBLAST analyses between *Si*, *Hpa*, *Psy* and *Gor* effector sequences. **B** PCA describing variance between sequence homology scores for each effector from Si, *Hpa*, *Psy* and *Gor*. **C** Venn diagram showing overlap in protein domains for microbial effectors predicted using EBI InterPro.**D** Absolute frequency of the most prevalent protein domains predicted for microbial effectors. **E** Network graph displaying interactions between effectors of *Si*, *Gor* and *Hpa* within the 12k search space. Effectors (hexagons) are classified based on the presence of an intrinsically disordered domain. Convergent host targets (circles) interact with both IDD and IDD-absent effector proteins.

**Supplemental figure 4.**
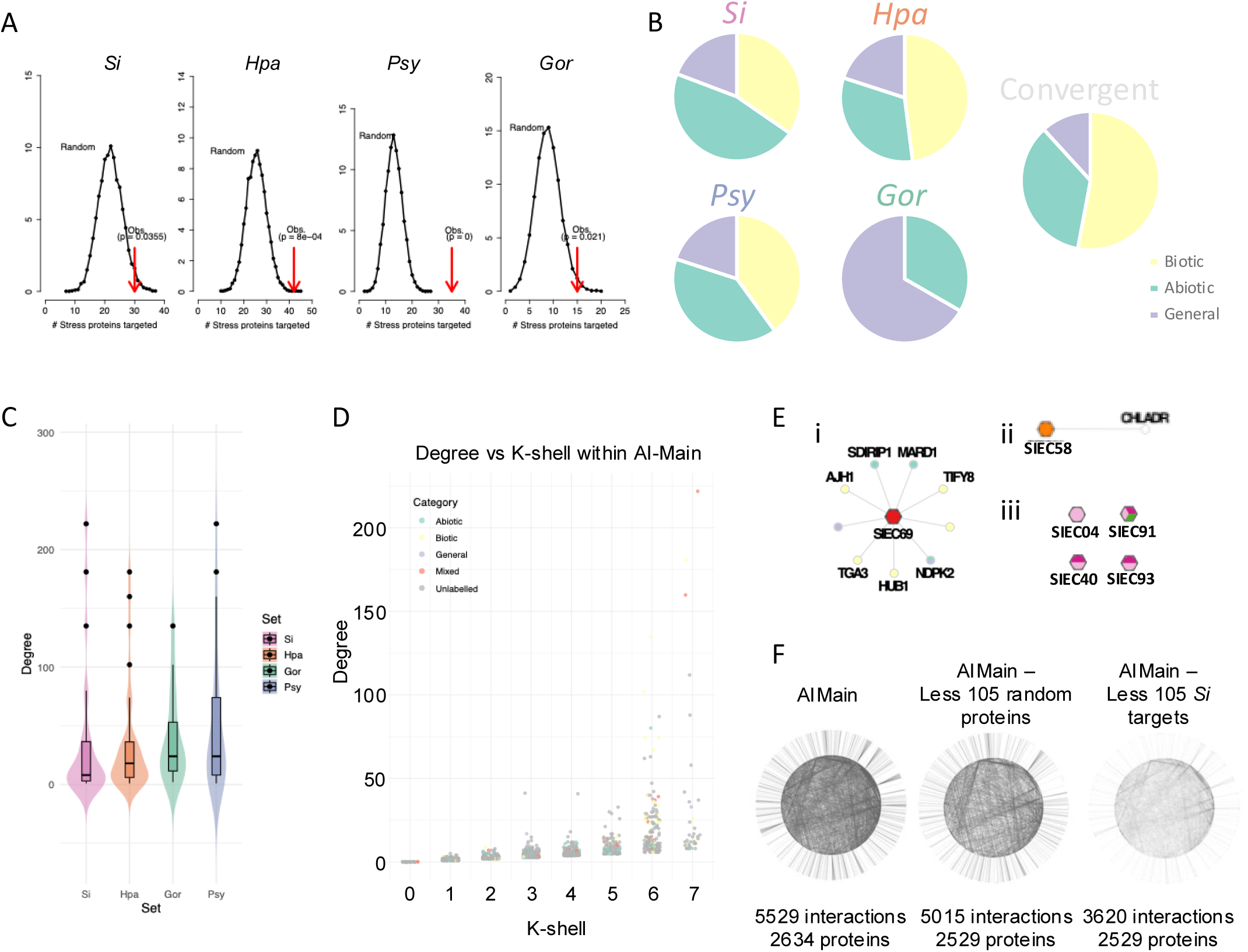
Effector proteins target stress responsive proteins. **A** Frequency of stress protein targeting by effectors in random networks (black dots) vs the observed rate of stress protein targeting (red arrows) in respective interactomes for *Si*, *Hpa*, *Psy* and *Gor*. **B** Categorisation of stress associated proteins amongst unique effector targets for *Si*, *Hpa*, *Psy Gor* and convergent targets. Charts comprise every semantic appearance of each stress category, i.e. a protein with a function in both abiotic and biotic stress appears in both sections of the pie chart. **C** Violin plot displaying the degree of effector targets within AI-Main sorted by organism. **D** Distribution of nodes within AI-Main when considering degree vs K-shell. Stress annotated proteins appear within each K-shell with no trend toward lower vs higher degree. **E** Subnetworks displaying interactions of SIECs used in abiotic stress phenotyping; (i) SIEC with multiple stress targets, (ii) SIEC with one stress related target, (iii) SIEC with known stress marker effects with no identified interactors. **F** Network graph displaying the interconnectivity of AI-Main. Removal of *Si* target proteins reduces the total number of interactions by ∼2,000, whereas the removal of an equivalent number of random proteins reduces the connectivity by ∼500.

**Supplementary Figure 5.**
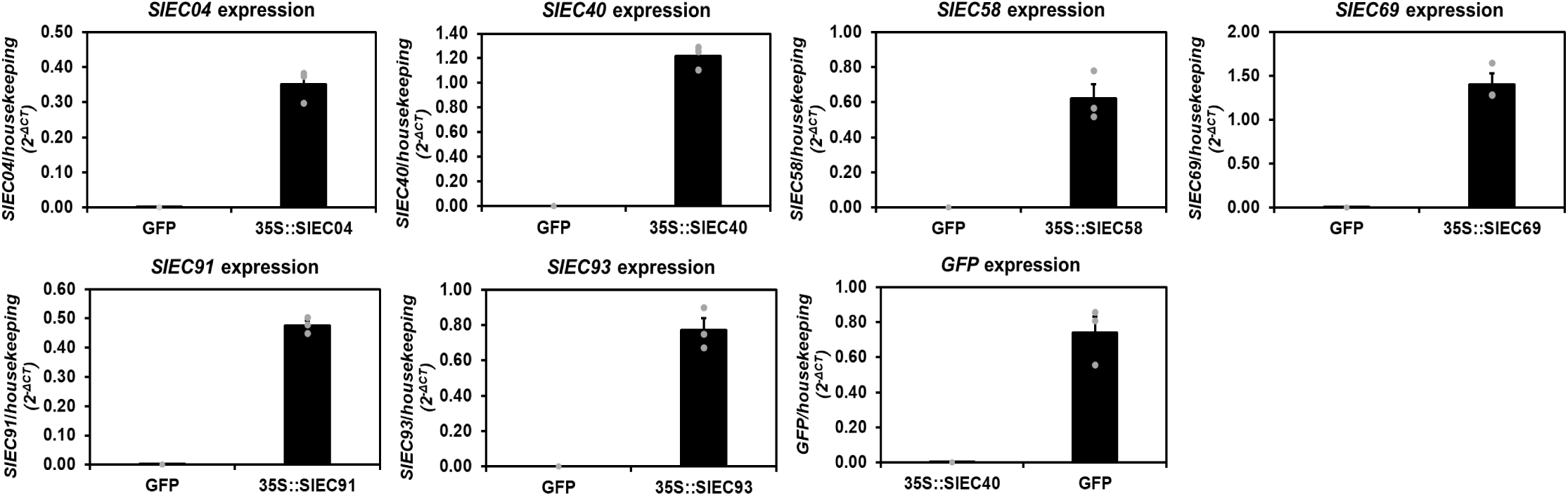
qRT-PCR-based quantification of *SIEC* expression in *SIEC*-expressing Arabidopsis seedlings. RNA of roots of 14-day old seedlings was used to analyse *SIEC* expression by qRT-PCR. *SIEC*-specific primers were used to confirm expression in *p35S::SIEC* lines relative to housekeeping genes *UBQ5* and *EF1*α. GFP-expressing plants served as negative control. Data shown is based on three technical replicates of RNA extracted from a pool of *p35S::SIEC* seedlings.

**Supplementary Figure 6.**
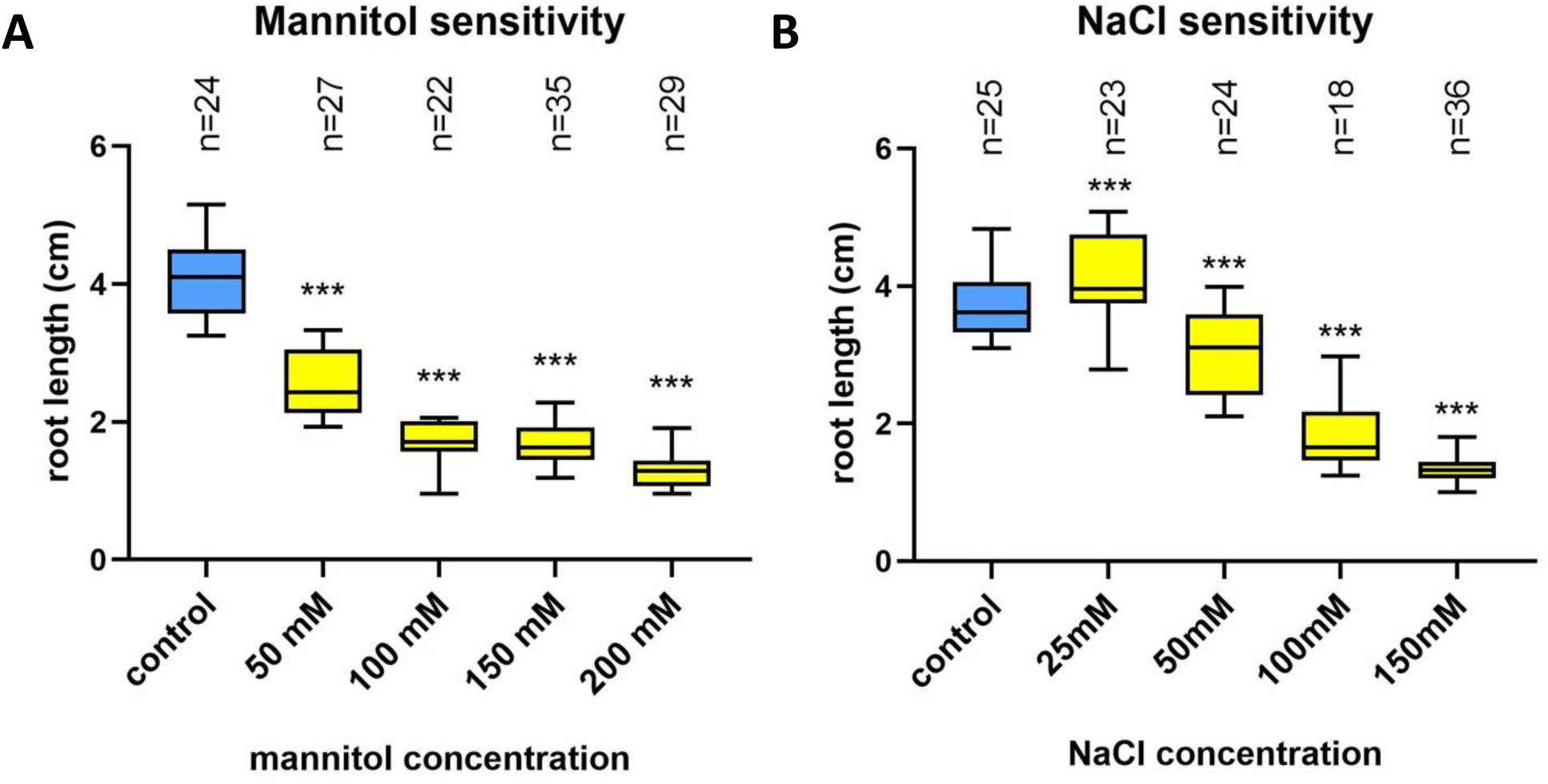
Mannitol and NaCl sensitivity assays. **A** Effect of ascending concentrations of mannitol and **B** NaCl on primary root length of 14-day old seedlings. Root length of control plants is indicated in blue. (***) p < 0.001; statistical significance was evaluated by ANOVA. Sample size (n) is indicated above each box.

**Supplementary Figure 7.**
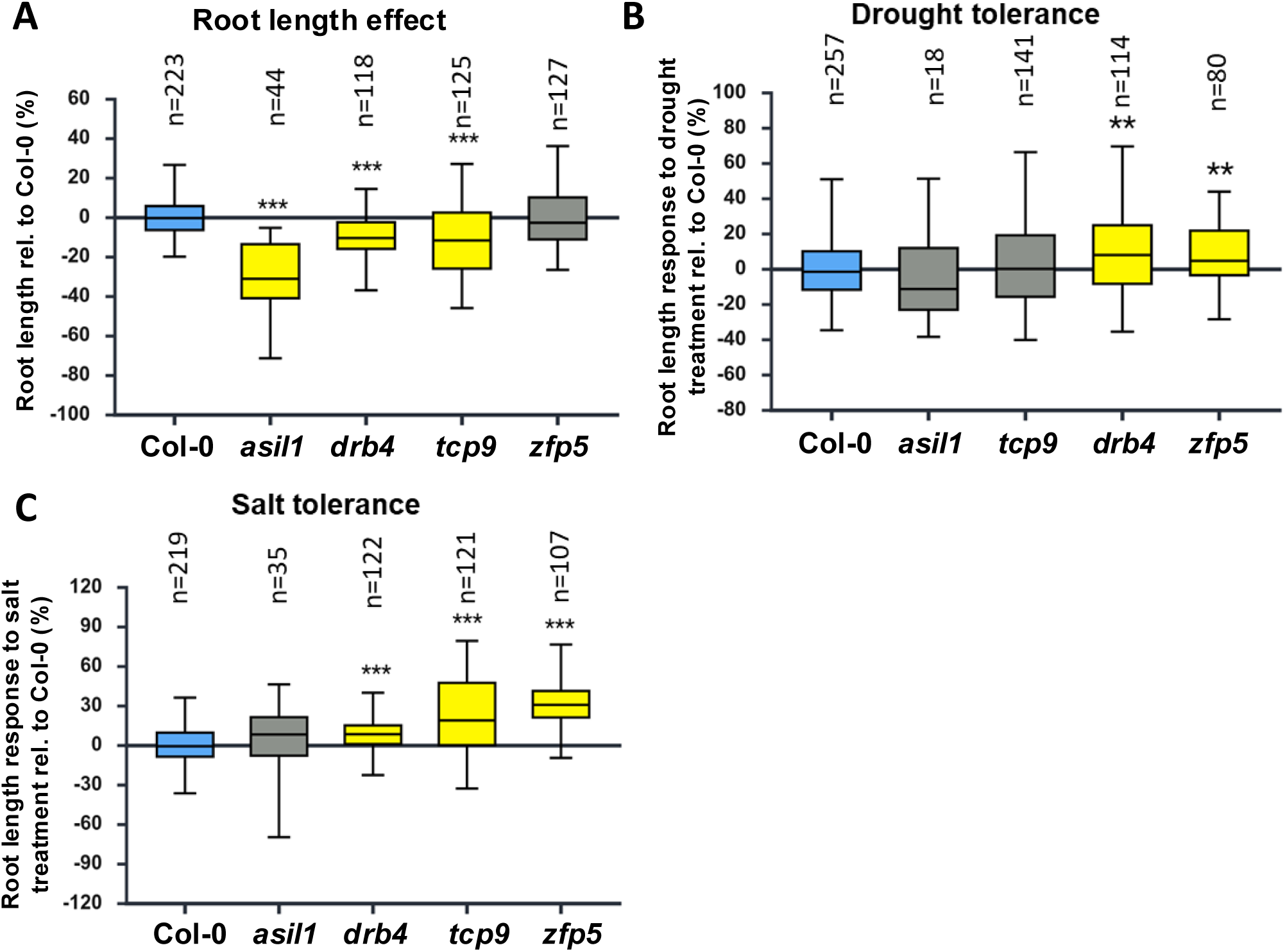
Phenotyping of target knockout lines. **A** Primary root length, **B** relative drought stress and **C** relative salt stress response of primary root length of 14-day old seedlings relative to Col-0 (%). Root length of control plants is indicated in blue. (***) p < 0.001; statistical significance was evaluated by ANOVA. Sample size (n) is indicated above each box.

## Supplementary Tables

**Supplementary Table 1. SIEC function in protoplasts.**

**Supplementary Table 2. SIEC Interactome and Target GO classifications.**

**Supplementary Table 3. Effector domain and sequence homology.**

**Supplementary Table 4. Functional domain analysis of microbial effectors.**

**Supplementary Table 5. Phenotyping of *Arabidopsis thaliana* mutants.**

**Supplementary Table 6. qRT-PCR raw data.**

**Supplementary Table 7. Primers used in this study.**

**Supplementary Table 8. Pathogen infection assay data.**

**Supplementary Table 9. SIEC and target cellular localisation.**

